# Multi-Omic Profiling Reveals the Opposing Forces of Excess Dietary Sugar and Fat on Liver Mitochondria Protein Acetylation and Succinylation

**DOI:** 10.1101/263426

**Authors:** Jesse G. Meyer, Samir Softic, Nathan Basisty, Matthew J. Rardin, Eric Verdin, Bradford W. Gibson, Olga Ilkayeva, Christopher B. Newgard, C. Ronald Kahn, Birgit Schilling

## Abstract

Dietary macronutrient composition alters metabolism through several mechanisms, including post-translational modification (PTM) of proteins. To connect diet and molecular changes, here we performed short- and long-term feeding of mice with standard chow diet (SCD) and high-fat diet (HFD), with or without glucose or fructose supplementation, and quantified liver metabolites, 861 proteins, and 1,815 protein level-corrected mitochondrial acetylation and succinylation sites. Nearly half the acylation sites were altered by at least one diet; nutrient-specific changes in protein acylation sometimes encompass entire pathways. Although acetyl-CoA is an intermediate in both sugar and fat metabolism, acetyl-CoA had a dichotomous fate depending on its source; chronic feeding of dietary sugars induced protein hyperacetylation, whereas the same duration of HFD did not. Instead, HFD resulted in citrate accumulation, anaplerotic metabolism of amino acids, and protein hypo-succinylation. Together, our results demonstrate novel connections between dietary macronutrients, protein post-translational modifications, and regulation of fuel selection in liver.

**Graphical Abstract:** Graphical Abstract

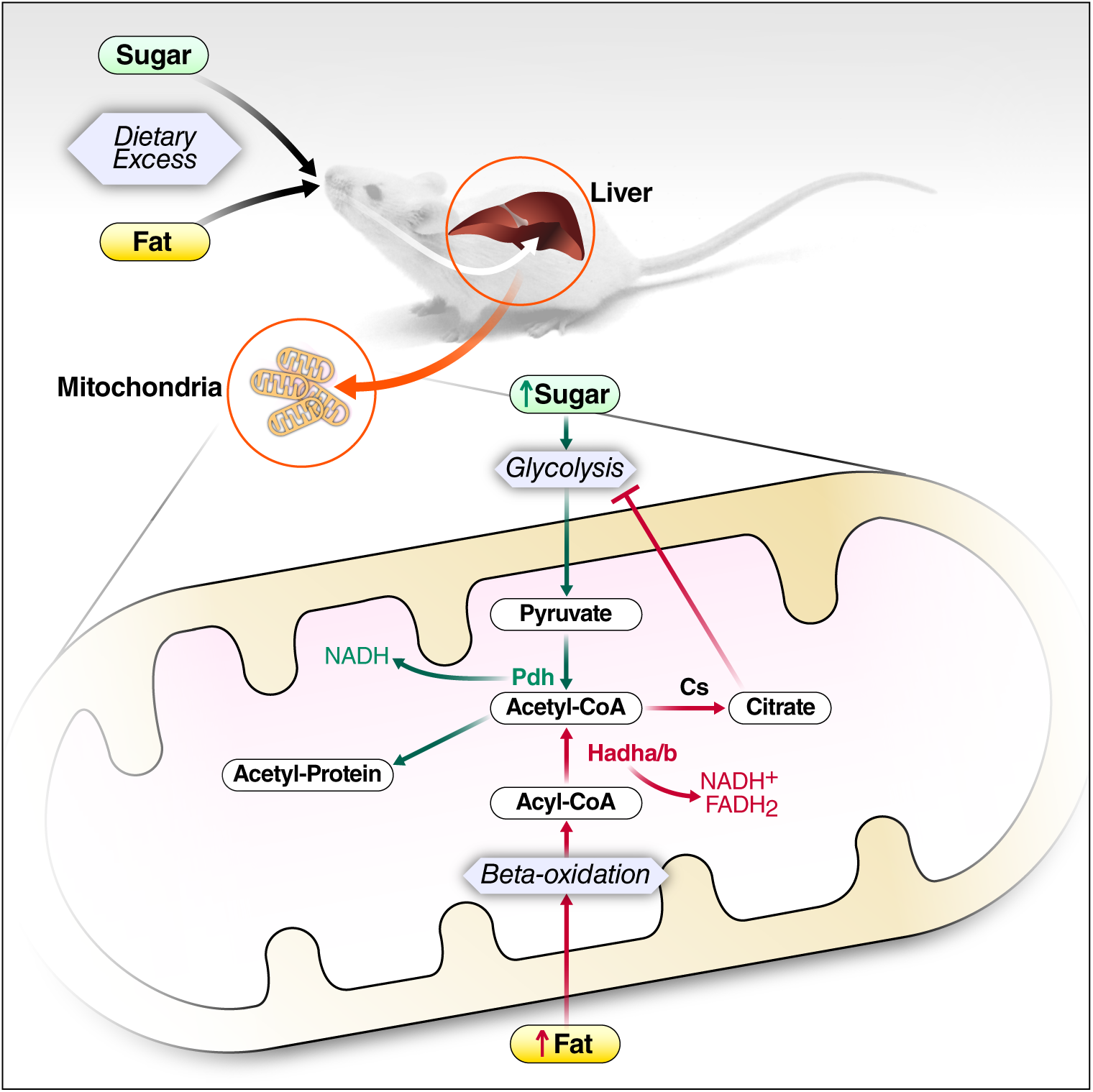

## Introduction

Lysine residues in proteins are a point of convergence for multiple PTMs, especially acylation. Lysine acetylation was first reported in nuclear proteins (Saxholm et al., 1982). More recently, immuno-affinity enrichment and mass spectrometry were used to discover that acetylation is also highly abundant in mitochondrial proteins (Kim et al., 2006). Modification of protein lysine residues by other acyl groups was discovered later, including succinylation (Zhang et al., 2011). The diversity of lysine acylations is mirrored by the diversity of reactive acyl-CoA (RAC) species, and includes malonylation and glutarylation (Anderson et al., 2017; Wagner and Hirschey, 2014). Similar to acetylation, succinylation extensively modifies mitochondrial proteins (Rardin et al., 2013a), but is also found in other cellular compartments (Weinert et al., 2013). Protein acetylation and succinylation are reversed by sirtuins, an NAD+-dependent class of lysine deacetylases (Bheda et al., 2016). In the mitochondria, Sirt3 and Sirt5 efficiently remove acetyl and succinyl groups, respectively (Rardin et al., 2013b; Nishida et al., 2015; Rardin et al., 2013a). Reversible modification of mitochondrial protein by acylation may allow tuning of mitochondrial function in response to nutrient availability, which costs less energy than degradation and synthesis of new proteins. Although there is a clear correlation between protein acylation and various phenotypes, the underlying causal relationship between phenotype, protein function, and protein acylation is still poorly understood.

Due to the prevalence of acetylation and succinylation on metabolic enzymes in the mitochondria, and the lack of evidence for a mitochondria-targeted acyl transferase, the growing consensus is that mitochondrial protein acylation is a non-enzymatic consequence of metabolism (Ghanta et al., 2013). Such non-enzymatic modification may be facilitated by a combination of high pH in the mitochondrial matrix, altered lysine side-chain pKa, the composition of nearby amino acids, and changes in levels of RAC under different metabolic conditions (Baeza et al., 2015; James et al., 2017). RAC composed of CoA and dicarboxylic acids appear to have a unique reactivity due to intramolecular base catalysis that can spontaneously form high-reactivity anhydrides *in vitro* and *in vivo* (Wagner et al., 2017). The non-enzymatic modification model is further supported by the correlation of RAC concentrations with the prevalence of specific RAC modifications, as well as the sirtuin deacetylase enzymes that remove them, across various tissues (Sadhukhan et al., 2016). Therefore, alteration of protein acylation should be interpreted in the context of RAC abundance and metabolism.

Mitochondrial protein hyper-acetylation has been associated with changes in protein function and with various genotypes and metabolic stressors, including Sirt3 knockout (Sol et al., 2012), ethanol exposure (Picklo, 2008), chronic caloric restriction (Schwer et al., 2009), ketogenic diet (Newman et al., 2017), or chronic HFD (Hirschey et al., 2010). HFD-induced mitochondrial protein hyper-acetylation has been studied extensively, and one study suggested protein acetylation is solely driven by acetyl-CoA produced from fatty acid oxidation (Pougovkina et al., 2014). Notably, mice fed HFD for 13 weeks, or Sirt3 KO mice, show accelerated development of metabolic disorders and liver inflammation (Hirschey et al., 2011). Compared to the number of studies that demonstrate hyper-acetylation, only a few conditions are associated with protein hyper-succinylation, such as mutations in isocitrate dehydrogenase associated with cancer (Li et al., 2015), succinyl-CoA ligase deficiency (Weinert et al., 2013), and knockdown of Sirt5 (Park et al., 2013; Rardin et al., 2013a).

Several reports describe how site-specific protein acylation can alter functions of proteins in multiple pathways. Similar to other protein modifications such as phosphorylation, protein acylation has been reported to cause both enzyme activation and inhibition. For example, carbamoyl phosphate synthase-1 (Cps1), which is required for ammonia clearance and amino acid catabolism, is deacetylated by Sirt5 to cause increased activity, thereby enabling the shift to amino acid catabolism during fasting (Nakagawa et al., 2009). Ketone body production is decreased in the absence of Sirt5, and an enzyme required for ketone body production, HMG CoA synthase 2 (Hmgcs2), appears to be inhibited by succinylation at K83 and K310 (Rardin et al., 2013a). In addition, hydroxyacyl CoA dehydrogenase (Hadha), a subunit of the trifunctional enzyme required for beta-oxidation of fatty acids, is inhibited by succinylation of K351 (Sadhukhan et al., 2016), and long-chain acyl-CoA dehydrogenase (LCAD) activity is inhibited by acetylation of K318 and K322 (Bharathi et al., 2013). Although less common, there are reports of functional activation of proteins by acylation. For example, the transcription factor ChREBP shows enhanced activity when acetylated on K672 (Bricambert et al., 2010). Thus, system-wide analysis of changes in protein acylation patterns, instead of effects on individual proteins, may be a roadmap for better understanding of regulatory mechanisms that drive physiologic phenotypes.

Although there are examples of acylation influencing metabolism, it is still unclear how diet composition can influence protein acylation. Since multiple acyl groups can modify any given lysine, studies that simultaneously assess the dynamic interplay between these modifications are needed to provide a more complete understanding of acylation and regulation of protein activity (Zhu et al., 2016). Here, we fed mice SCD or HFD with or without supplemented fructose or glucose, and quantified liver mitochondrial proteins, mitochondrial protein acetylation and succinlyation, as well as hepatic organic acids, acyl CoA, acylcarnitine, and amino acids levels. To rapidly and comprehensively quantify acetylation and succinylation, we combined immunoprecipitation, quantitative, label-free, data-independent acquisition (DIA) mass spectrometry (Collins et al., 2017; Gillet et al., 2012), and automated data analysis with the PTM identification and quantification from exclusively DIA (PIQED) workflow (Meyer et al., 2017). This multi-omics approach, which combines quantitative analysis of proteins, PTMs, and metabolites, provides a comprehensive view of molecular responses to chronic changes in diet composition, and resultant changes in macronutrient signaling and selection in the liver. We find a dichotomous response to excess dietary sugar and fat; feeding excess glucose for 10 weeks (and to a lesser extent fructose) induces mitochondrial protein hyper-acetylation, but HFD alone does not. Instead, chronic HFD caused protein hypo-succinylation and increased citrate, a known inhibitor of glycolysis.

## Results

### Overview of Global Diet-Induced Remodeling of Mouse Liver Proteins and Protein Acylation

To study how individual macronutrients can contribute to metabolic outcomes, cohorts of mice were fed six different diets. Mice fed SCD or HFD were provided regular water, or water supplemented with 30% fructose or 30% glucose. The diets were administered for 2 or 10 weeks to represent relatively acute and more chronic effects of dietary exposure (**Figure 1A**). The diet groups are abbreviated as: SCD plus water (control), SCD plus fructose (F), SCD plus glucose (G), HFD plus water (H), HFD plus fructose (HFD+F), and HFD plus glucose (HFD+G). Liver tissue was used to measure a panel of organic and amino acids, as well as acyl CoA and acylcarnitine metabolites by targeted mass spectrometry. In parallel, mitochondria were isolated from liver tissue in order to measure the liver mitochondrial proteome, and protein lysine acetylation and succinylation (**Figure 1A**).

**Figure 1:**
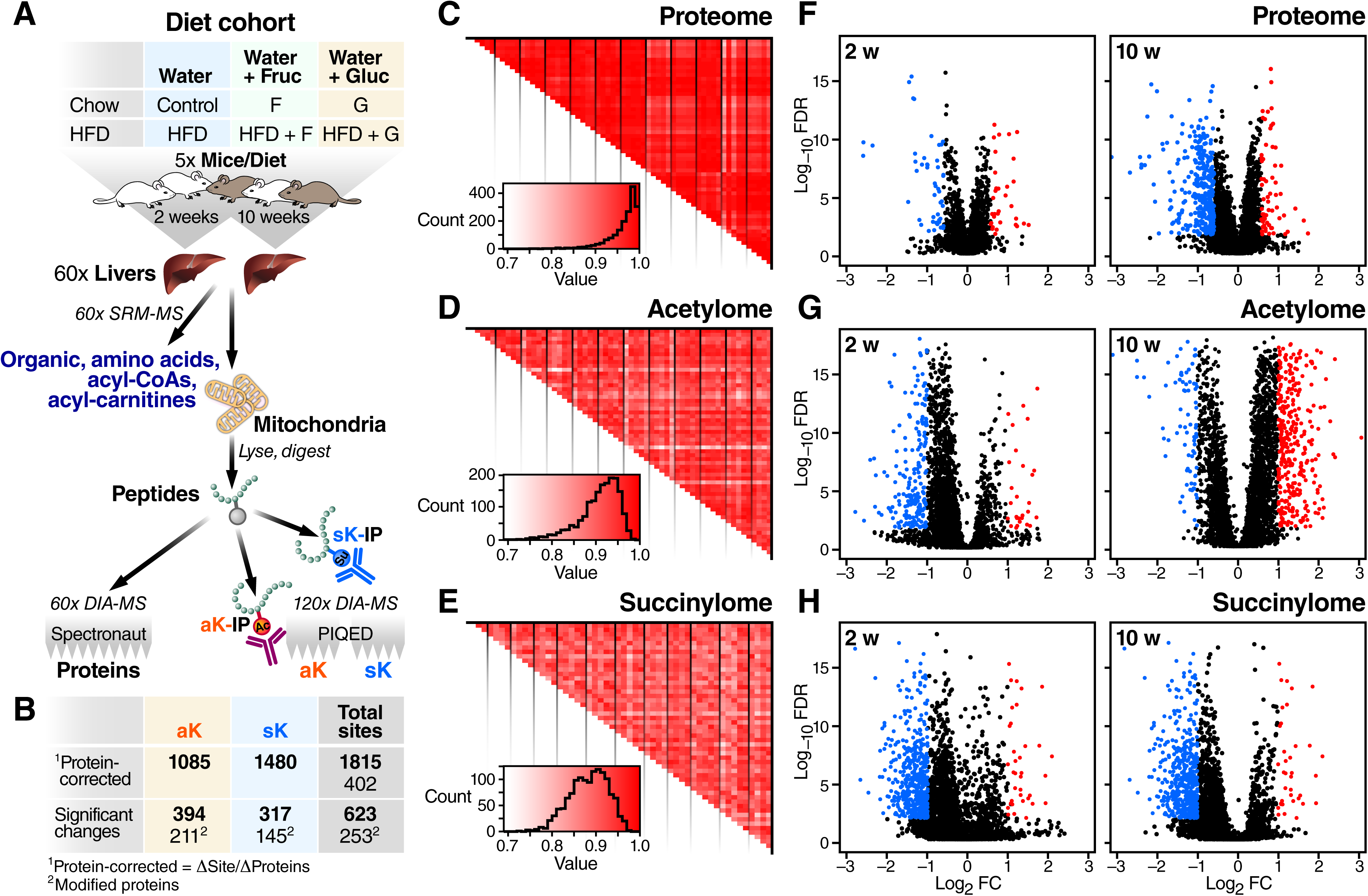
Diet-induced changes in mouse liver proteome, acetylome, and succinylome. **a,** Schematic of the dietary conditions, sample preparation, and analysis workflow. Five mice from each dietary cohort were sacrificed and a portion of liver tissue was used for quantification of organic and amino acids. Mitochondria were also isolated from separated portions of liver tissue and used for measuring alterations in abundance of proteins, protein acetylation, and protein succinylation. **b,** Calculation of protein-normalized acylation and overview of the qualitative and quantitative results. **c-e,** Pearson correlations comparing all peak areas of each biological replicate with all peak areas of other biological replicates, including those from other diet cohorts, for each of the following datasets: proteome **(c),** acetylome **(d),** and succinylome **(e)**. The vertical black lines delineate diet groups. **f-h,** Volcano plots showing magnitude and significance of all measurements compared to the 2-week or 10-week control cohort for the proteome **(f),** acetylome **(g),** and succinylome **(h).**

Using recent mass spectrometry advances (see Methods) we identified 862 proteins, 1,465 acetylated peptides and 1,957 succinylated peptides. Measurement of both protein and lysine modification levels from the same samples allowed correction of acylation changes by changes in protein expression, which was automatically performed by PIQED (Meyer et al., 2017). This correction, which required protein-level quantification for all acylation site-level quantification, reduced the number of protein abundance-corrected acetylation and succinylation sites to 1,085 and 1,480, respectively. A total of 1,815 modified lysine residues on 402 proteins (**Figure 1B**) were identified and quantified. Among those, 750 lysine residues were confidently identified as both acetylated and succinylated **(Figure S1)**.

Quantified proteins and protein acylation sites from each condition showed high reproducibility among all biological replicates, with the average correlation coefficients (Pearson r) of 0.96, 0.91, and 0.88, for proteome, acetylome, and succinylome measurements, respectively (**Figure 1C-E)**. Our three datasets are therefore of similarly high quality and reproducibility as a recent large-scale phospho-proteomics resource study (Robles et al., 2017). We compared each diet to the corresponding 2- or 10-week control group and found that 623 lysine sites in 253 proteins showed at least one statistically-significant diet-induced change (fold change > 2, FDR <0.01, **Figure 1B, 1G, 1H**). The overall trends in changes in protein abundance, protein acetylation, and protein succinylation are evident from volcano plots of -log_10_(FDR) versus log_2_(FC) (**Figures 1F-H**).

### Chronic HFD and Glucose Induce Distinct Remodeling of Mouse Liver Mitochondrial Proteins

Automatic correction of acylation site quantification by protein abundance performed by PIQED software was critical in this study due to significant diet-induced mitochondrial proteome abundance changes; 276 proteins significantly changed in at least one comparison (fold change >1.5, q-value < 0.01, **Figure 1F, Figure 2A**). Most of the protein-level changes resulted from 10 weeks of feeding the three HFDs. Because the liver serves as an important regulator of glucose and lipid homeostasis (Frayn and Kingman, 1995), we also compared the 10-week G group with the 10-week HFD group (last column in **Figure 2A**). Proteins with significantly altered abundance in the 10-week glucose group compared to the 10-week HFD group clearly segregate into separate functional groups **(Figure 2B)**. Mice fed HFD for 10 weeks showed higher abundance of mitochondrial proteins involved in amino acid catabolism than mice fed excess glucose for 10 weeks, whereas mice fed glucose for 10 weeks had increased levels of several proteins involved in lipid metabolism (e.g. Ehhadha and several cytochrome P450s) and cellular redox homeostasis (e.g. Gstm1, Gsta3, Gstp1, and SOD1). The increased levels of redox proteins are consistent with our finding that mice fed excess glucose produce less reactive oxygen species (ROS; see companion paper), and that some organisms use sugar metabolism to reduce oxidative damage (Levin et al., 2017). Additionally, acid phosphatase 5 (ACP5), which degrades FMN to riboflavin, is lower in HFD, consistent with the increased need for the FAD cofactor in fatty acid oxidation.

**Figure 2:**
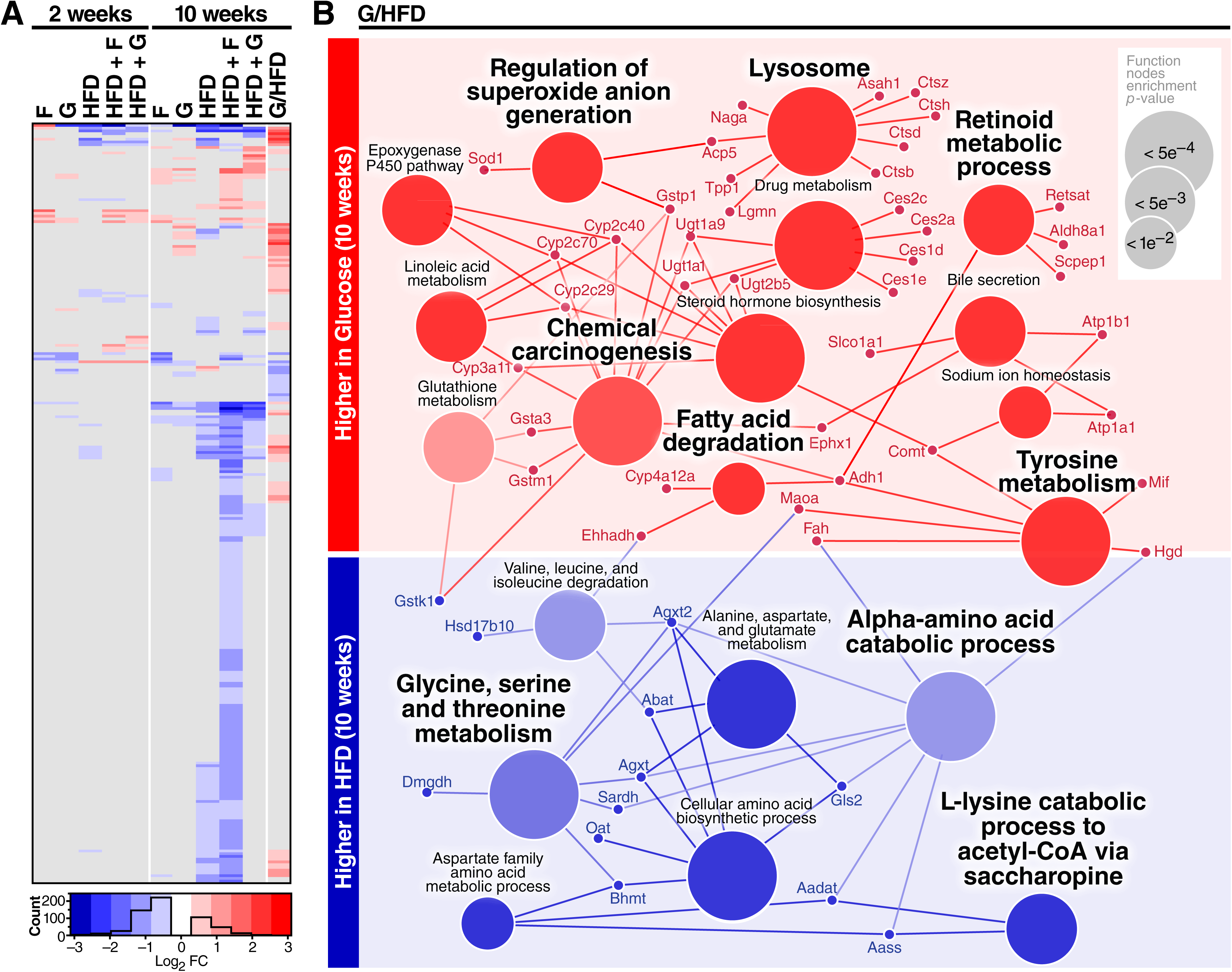
Quantitative protein-level results overview. **a,** Heatmap of 276 proteins that show at least one statistically-significant diet-induced change (q-value < 0.05, fold-change > 1.5). Values in grey are not significant at this threshold. Each diet is compared to the appropriate 2- or 10-week control, except the last column shows the direct comparison of 10-week glucose divided by 10-week HFD. **b**, Proteins showing altered abundance (small circles) between the 10-week glucose cohort and the 10-week HFD cohort in the last column of **(a)** were analyzed for GO and KEGG ontology term enrichment (large circles). Blue indicates lower abundance in 10-week glucose group compared to 10-week HFD group (or higher abundance in the 10-week HFD group than the 10-week Glucose group) and red indicates higher abundance in the 10-week glucose / 10-week HFD comparison (or lower abundance in the 10-week HFD group than the 10-week glucose group).

Among proteins that increased in the glucose-fed group compared to the HFD group were several enzymes involved in drug metabolism and lipid hormone metabolism, such as cytochrome p450s. Our data suggest that the less deleterious metabolic impact of excess glucose ingestion compared to excess HFD ingestion (Softic et al., 2017) may be mediated in part by glucose-induced increases in abundance of proteins that perform arachidonic acid hydroxylation (Cyp2c29, Cyp2c40, Cyp2c70), a process that can either activate or inactivate the inflammatory response, and in the case of Cyp2c29, produces an anti-inflammatory epoxyeicosatrienoic acid (EET) (Thomson et al., 2012). Also, Cyp3a11, a protein that hydroxylates various substrates including testosterone, was higher in glucose-fed animals than HFD-fed animals after 10 weeks. These protein-level changes are consistent with studies that show obesity-related reduction of redox capacity, altered drug metabolism, and increased inflammation in obese mice and humans (Merrell and Cherrington, 2011; Park et al., 2010).

### Diet-Induced Changes in Amino Acids, TCA Intermediates, Acyl-CoAs, and Acyl-carnitines

The dichotomous changes observed in protein abundances due to chronic excess ingestion of sugar compared to chronic excess ingestion of fat are reflected by the metabolite changes observed in livers from the 10-week cohorts. In agreement with the increased levels of several proteins involved in amino acid catabolism, we found a corresponding decrease in multiple amino acids in livers from the HFD groups. Relative to the 10-week control group, mice fed any HFD exhibited significant decreases in glycine, serine, leucine/isoleucine, glutamate/glutamine, methionine, histidine, phenylalanine, tyrosine, and the urea cycle intermediates ornithine and citrulline **(Figure 3A)**. Interestingly, feeding fructose for 10 weeks had no effect on amino acid levels compared to the control group. Unique to the chronic glucose-fed group, levels of arginine, glutamate/glutamine, and serine were increased relative to the 10-week controls. Chronic dietary glucose supplementation was also associated with more modest decreases in methionine and valine. The only other condition where valine was decreased was 10-week HFD+G, suggesting that excess dietary glucose potentiates valine catabolism in the HFD setting. These diet-specific changes were not evident at the 2-week time point, and are thus considered as chronic and not acute responses to the various diets.

**Figure 3:**
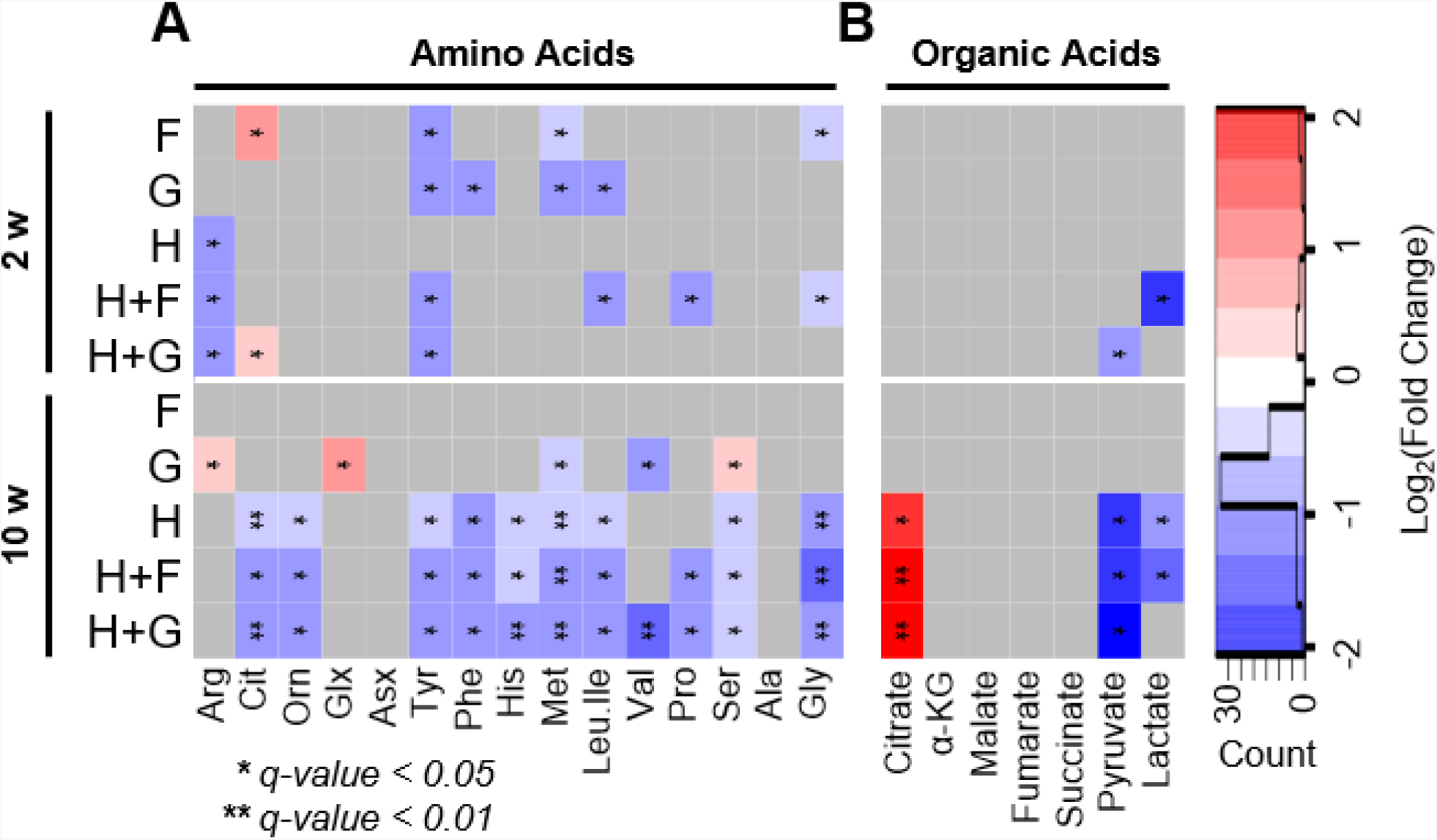
Diet-Induced changes of organic acid and amino acid abundances. Heatmaps show all statistically significant (any fold change, * q-value < 0.05, ** q-value < 0.01) abundance changes in the liver tissue relative to the appropriate 2-week or 10-week control group. **a,** Amino acids and, **b,** organic acids. n=5 mice.

Since amino acids can serve as a source of anaplerotic substrates for replenishment of the tricarboxylic acid (TCA) cycle, we also quantified TCA cycle intermediates **(Figure 3B)**. In liver tissue from mice fed HFD, HFD+F, or HFD+G for 10 weeks, citrate was significantly elevated and pyruvate was sharply decreased. Lactate was also lowered in the HFD and HFD+F groups. Citrate is known to inhibit glycolysis in all animals except insects (Newsholme et al., 1977), which reinforces the selection of fat as the primary fuel source. This increase in citrate was not evident after 2 weeks of diet feeding, whereas lactate was decreased after 2 weeks of feeding of HFD+F and pyruvate was decreased after 2 weeks of feeding of HFD+G, again suggesting a time-dependent impact of the diets on metabolic fuel selection.

Given the apparent connection between excess dietary fat intake and increased amino acid catabolism, levels of various hepatic acyl-CoA and acyl-carnitines were compared to the respective 2-week or 10-week controls (**Figure S2**). Livers from mice fed HFD or HFD+F accumulated various medium and long-chain acyl-CoAs and acyl-carnitines, suggesting high rates of influx of lipids into the □-oxidation pathway and accumulation of incompletely oxidized intermediates. Strikingly, these intermediates did not accumulate in the chronic SCD plus glucose-fed group. Moreover, addition of glucose to HFD almost completely abrogated the accumulation, possibly by increasing lipid metabolism. This interpretation is supported by direct measurements of fatty acid oxidation, CPT1 activity and malonyl CoA levels, as reported in the companion paper. To be conservative, we cannot rule out the possibility that glucose acts upstream to prevent lipid utilization. Interestingly, acetyl-CoA and succinyl-CoA, which are thought to non-enzymatically induce protein modification, showed minimal alteration among the tested dietary conditions; acetyl-CoA was increased due to acute glucose feeding and decreased due to chronic HFD, while succinyl-CoA was only altered in chronic HFD+G, where it increased. The latter result also supports glucose-related increase in valine metabolism, as valine is metabolized to succinyl-CoA. We also note an increase in triply-unsaturated C10 acyl-CoA and acyl-carnitine that results from all HFDs.

### Opposite Changes in Protein Acetylation and Succinylation in Response to Chronic Dietary Sugar or Fat

Using the stringently-filtered set of site-level acylation changes relative to respective 2-week or 10-week controls, several general trends become apparent **(FDR <0.01, fold-change>2, Figure 4A)**. Both lysine acetylation and succinylation showed more changes after 10 weeks compared to 2 weeks of feeding, and overall acetylation changed more than succinylation. Most of the sites that exhibit decreased acylation due to any diet show less than 50% overlap with previously-described Sirt3- or Sirt5-regulated acylation sites (Hebert et al., 2013; Rardin et al., 2013b, 2013a) **(Figure 4B)**. The low overlap of diet-regulated sites and sirtuin-regulated sites suggests that either the sites were not identified in the previous studies, possibly due to the lack of dietary stress or abundance below the detection limit, or that these diet-regulated sites may not be regulated by the sirtuins. Similar to the divergent changes found in proteins and metabolites, dietary sugar and fat also have dichotomous effects on acetylation and succinylation changes with opposite trends for the two macronutrients for each PTM. Global hyperacetylation was observed in response to chronic glucose or fructose supplementation of chow-fed mice, but HFD was associated with a nearly equal number of increased and decreased sites. Interestingly, addition of glucose to HFD was sufficient to induce some protein hyperacetylation, while added fructose was not. In contrast, chronic feeding of either sugar had almost no effect on protein succinylation, whereas all three of the HFDs induced general hypo-succinylation; the 10-week HFD+F cohort showed the most dramatic effect.

**Figure 4:**
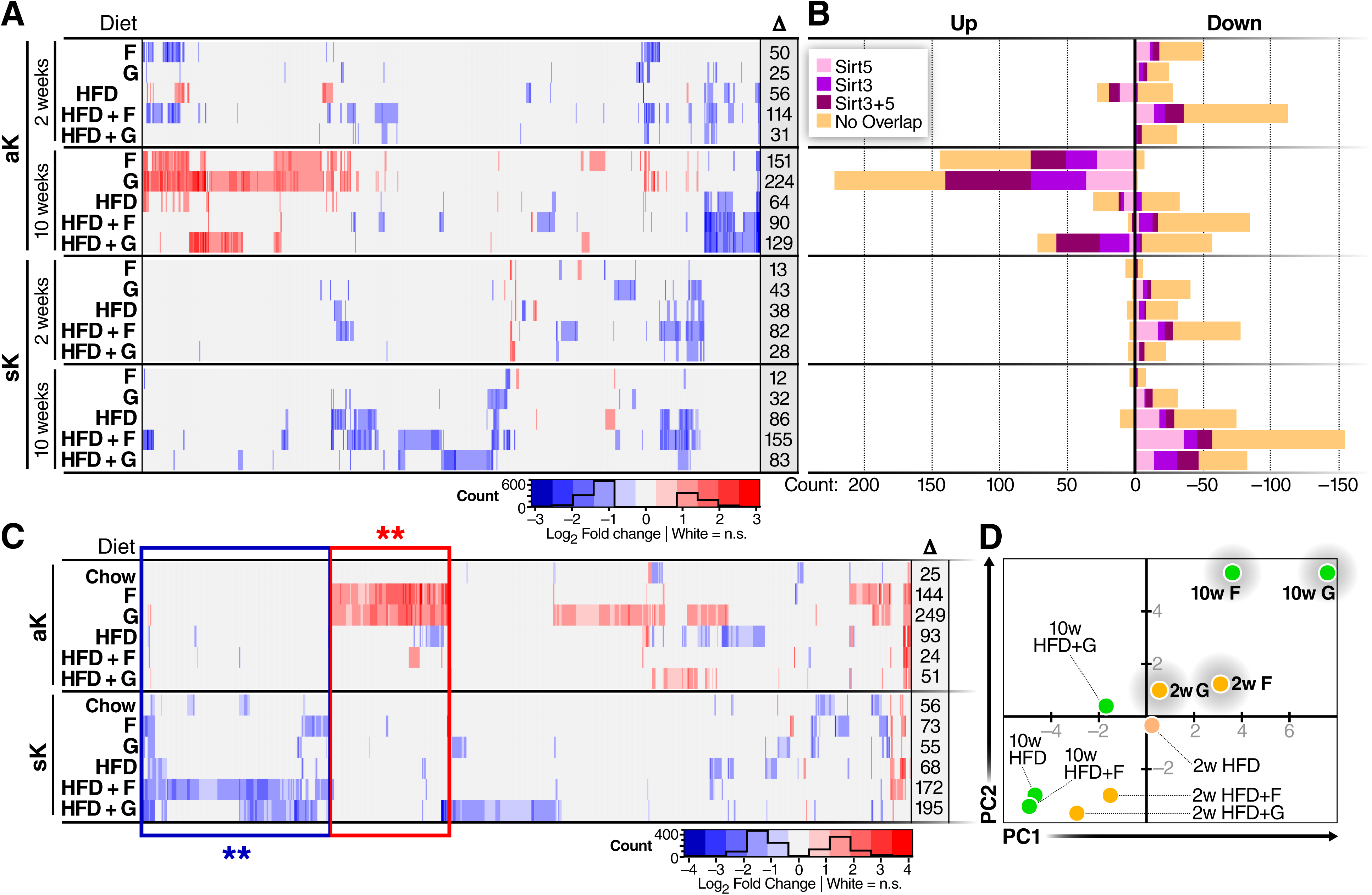
Diet-induced remodeling of liver mitochondria protein acetylation and succinylation. **a,** Heatmap showing 623 modified lysine residues from 253 proteins with at least one statistically-significant change in any diet (FDR<0.01, fold change >2). Columns show acylation sites across the diets, timepoints, and acylation type, and color indicates the fold change of that site relative to the appropriate 2-week or 10-week control. At the right of each row, the total number of significant changes are indicated. **b,** Number of lysine sites showing statistically-significant increase or decrease in abundance of acetylation or succinylation for each diet, and the overlap of those sites with previously-reported Sirt3 or Sirt5-regulated modification sites. **c,** The same as **(a)**, however, these heatmaps illustrate comparisons of 10-week modification abundance over the 2-week abundance. 638 sites in 244 proteins show a time-dependent change due to at least one diet. The numbers at the end of each row indicate the number of significant changes. Groups of boxed sites denote the sets used for functional analysis in figure 5. **d**, Principle component analysis of the acetylation site levels show segregation of the dietary sugars in chow background from all other groups.

Similar to the protein abundance and metabolite data, changes in lysine acylation were more prominent after the chronic, 10-week feeding compared to the acute 2-week diets. To understand this trend, 10-week acylation was compared to 2-week acylation quantities for each diet **(Figure 4C)**, instead of comparing with the respective 2-week or 10-week control groups as shown in figure 4A. Not surprisingly, minimal differences were observed comparing the 10-week controls to the 2-week control diets, importantly ruling out maturation or development as causes. Thus, protein acylation changes are dependent on both the type of nutrient excess and the length of exposure. The number of altered acetylation sites found in F, G, and HFD cohorts was similar in the time-dependent analysis (10-week / 2-week, same diet condition) as reported for the diet-dependent analysis (10-week diet /10-week control). However, the time-dependent analysis produced a more modest number of changes in acetylation in the HFD+F and HFD+G cohorts. In contrast, more altered succinylation sites in the F, G, and HFD+G diet groups were found using the time-dependent analysis (10-week / 2-week) compared to the diet-dependent analyses (10-week diet/10-week control). Many of the time-dependent changes were not found in the diet-dependent comparison; by including the time dependent changes, the number of sites with at least one diet-induced change increased by 38% to 861 sites across 301 proteins (**Figure S3A**). Notably, in half of the diets, hypo-succinylation was more time-dependent than hyper-acetylation, even though succinyl-CoA is more reactive than acetyl-CoA **(Figure 4C)** (Wagner et al., 2017).Together, this analysis of time-dependence indicates that sugar-dependent hyper-acetylation and fat-dependent hypo-succinylation increase with time, with the most significantly altered acetylation resulting from glucose supplementation and the most significantly altered succinylation profile occurring in response to HFD+F feeding.

To further probe the molecular details of diet-induced acylation, we assessed similarity among the dietary groups using both principle component analysis (PCA) and hierarchical clustering **(Figure 4D, Figure S3B)**. The PCA of acetylation changes distinctly segregates the sugar-fed groups from the HFD groups: all groups fed only sugar are in one quadrant and nearly all the HFD-fed groups are in another quadrant. The 10-week HFD+G group is at the opposite end of the PCA plot compared to the 10-week G group. Among the HFD groups, we obseved a switch in clustering from 2 to 10 weeks. The 2-week HFDs with added sugar cluster together while HFD without sugar is separate, but the 10-week HFD without sugar clusters closer to HFD+F, while HFD+G is less similar. PCA of diet groups based on succinylation changes also segregates the groups, and the segregation of the 2-week diets mirrored the acetylation changes, but the 10-week diets show sugar-dependent segregation **(Figure S3C)**. Notably, the 10-week HFD, HFD + F, and F groups cluster together. According to the unsupervised hierarchical clustering of diets and acetylation changes, the diets are related primarily by the presence of HFD, and secondarily by the presence of sugar. **(Figure S3B)**. Clustering of succinylation, in contrast, is primarily based on the length of the diet, and secondarily based on the presence of dietary glucose **(Figure S3C)**.

### Diet-Induced Acylation Changes Converge at the TCA Cycle and Amino Acid Catabolism

Although distinct clusters of acetylation and succinylation sites are evident from the heatmaps in figure 4A and 4C, a combined pathway enrichment analysis was done using the proteins containing sites enclosed by boxes marked with double asterisks in figure 4C (**Figure 5**). Twelve proteins in the electron transport chain (ETC) and eleven proteins in the combined TCA cycle/amino acid metabolism pathways showed sugar-dependent hyper-acetylation. In contrast, animals fed a HFD showed hypo-succinylation of seventeen proteins involved in amino acid metabolism and the TCA cycle, and eight proteins in the ETC (**Figure 5**). Several proteins involved in fatty-acid beta-oxidation were also found to be hyper-acetylated in response to sugar feeding and hypo-succinylated by HFDs. Eight proteins in the combined TCA cycle and amino acid metabolism pathways are commonly affected by feeding of sugar and HFD: Cs, Dlst, Idh2, Mdh2, Suclg2, Cps1, Got2, Hmgcs2. The acylation changes on these proteins follow the global trend of hyperacetylation in sugar supplemented diets and hypo-succinylation in HFD **(Figure S4)**.

**Figure 5:**
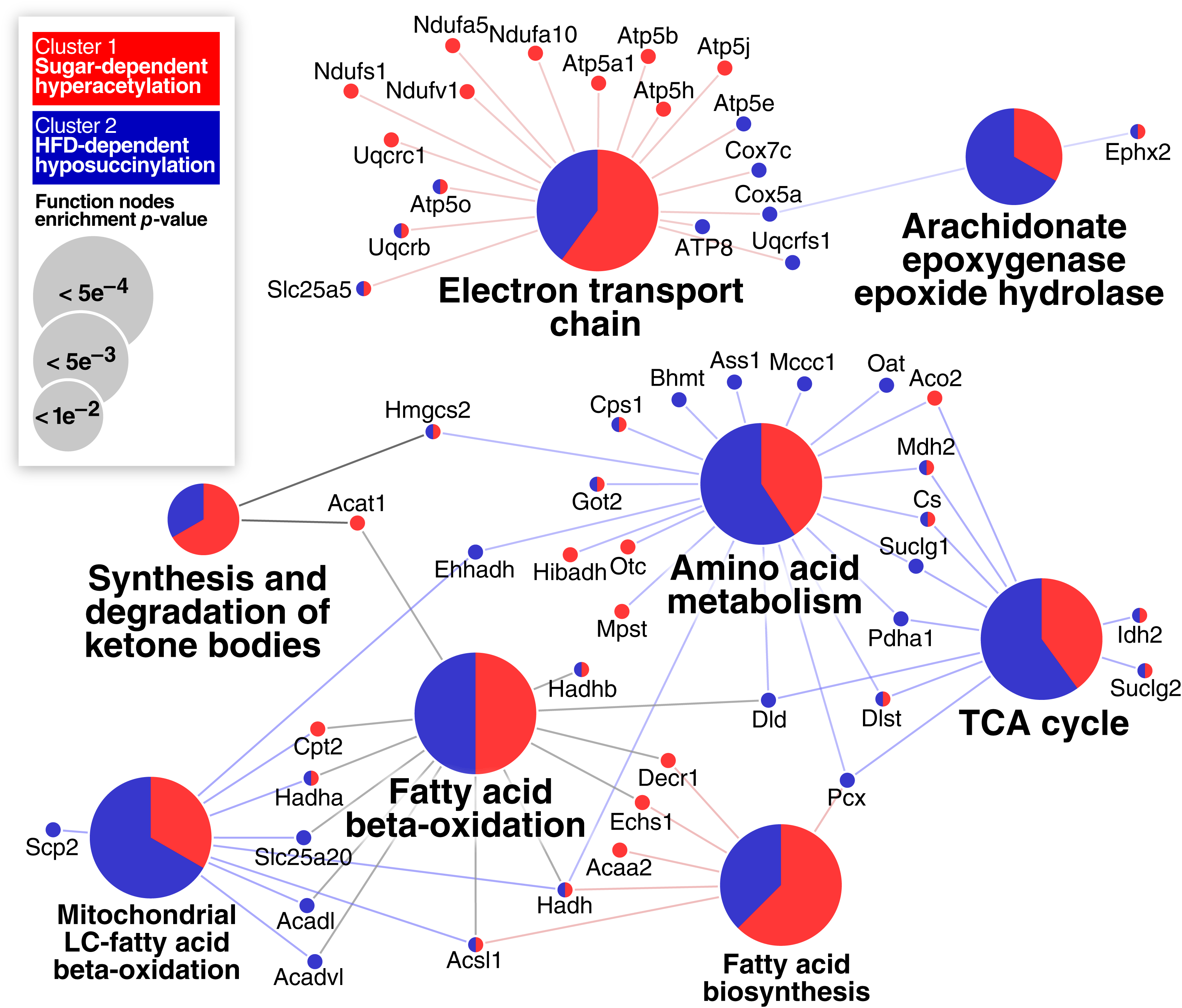
Functional grouping of the proteins showing dichotomous acylation responses. Enriched functional categories of proteins containing sites (denoted by boxes and asterisks in **Figure 4C**) that undergo site-specific glucose-induced hyper-acetylation (red) or HFD-induced hypo-succinylation (blue). The proportion of red and blue in the functional circles reflects the proportions of proteins from each group that map to this function.

Given the convergence of fat- and sugar-induced acylation changes on proteins in the TCA cycle, we examined the acetylation and succinylation changes on these proteins across all conditions in more detail **(Figure 6)**. Every protein in the TCA cycle with at least one quantified acetylation change featured glucose-dependent hyper-acetylation, and at least one HFD-dependent hypo-succinylation site. Among the enzymes with this pattern were pyruvate dehydrogenase, which generates acetyl-CoA from sugar, and the trifunctional enzyme, which is involved in generating acetyl CoA from fat. In the current study, HFD feeding increased citrate, and decreased multiple amino acids, as well as pyruvate and lactate. The decrease in lactate and pyruvate is consistent with the classical citrate-regulated decrease in glycolysis through inhibition of phosphofructokinase, whereas the decrease in amino acids may indicate their increased use as anaplerotic substrates to combine with fat-derived acetyl CoA for citrate production. HFD is known to cause a loss of metabolic flexibility, such that fatty acid oxidation becomes predominant in both the fasted and fed states (Galgani et al., 2008; Muoio, 2014), as opposed to animals fed on chow diets in which normal switching occurs between fatty acid oxidation in the fasted state and glucose oxidation in the fed state. Taken together, these findings suggest that HFD-dependent hypo-succinylation of multiple proteins may play a role in the adaptive response to HFD feeding, resulting in maximized oxidation of fat, suppression of glucose utilization, and increased amino acid catabolism to provide anaplerotic substrates.

**Figure 6:**
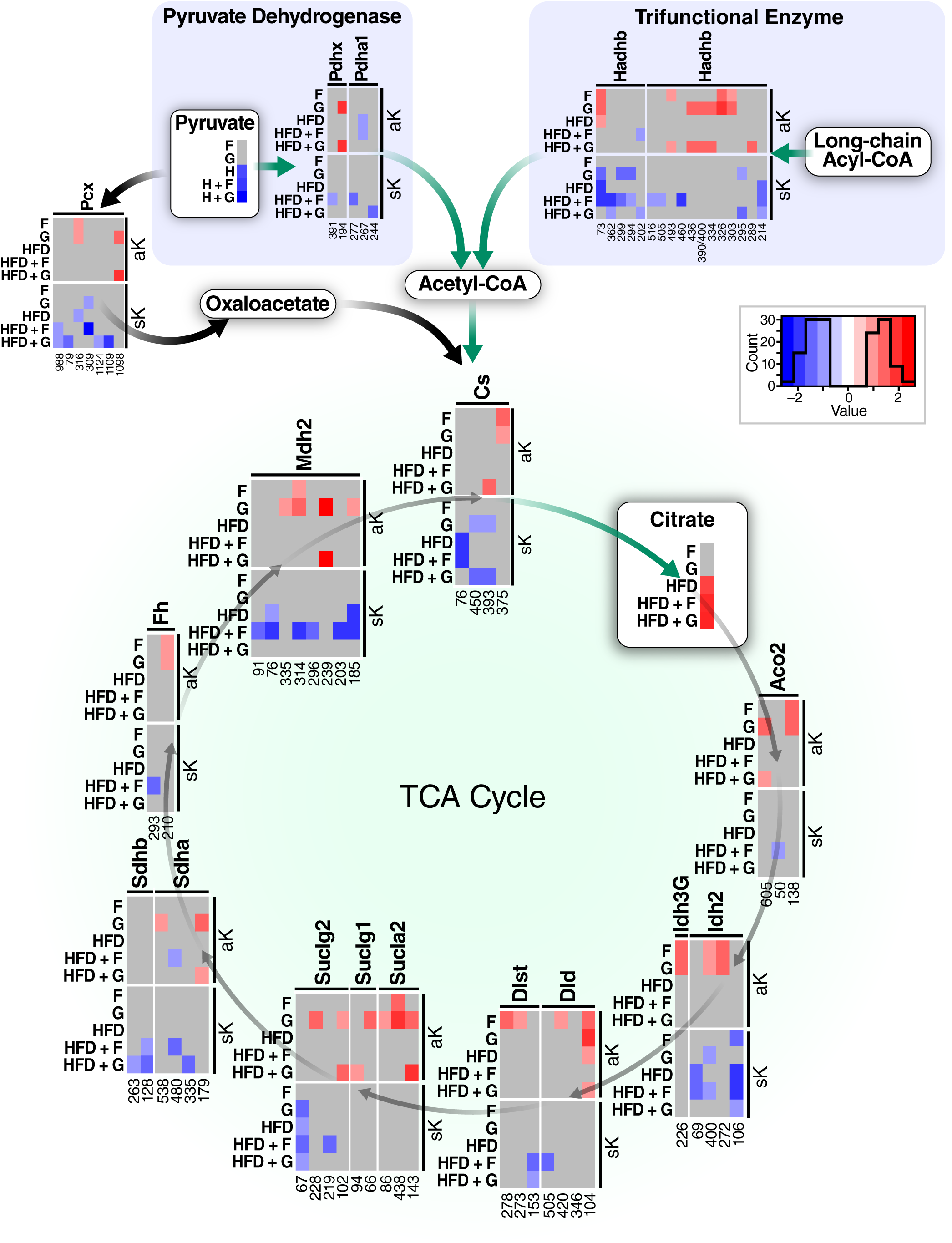
Acetylation and Succinylation changes on lysine residues from proteins in the tricarboxylic acid cycle and the proteins that generate acetyl-coA. Schematic of proteins involved in acetyl-CoA generation and the TCA cycle with at least one detected change in acylation after 10 weeks of feeding. Every detected protein in the TCA cycle has at least one glucose-dependent site of hyper-acetylation, and at least one HFD-dependent hypo-succinylation. Pyruvate and citrate quantification from **Figure 3** are also shown here.

Since all our data points to functional alteration of amino acid metabolism, acylation of proteins in the amino acid metabolism pathway were compared with those from the TCA cycle. Among the proteins in amino acid metabolism with altered acylation, a total of 116 sites in 27 proteins change due to at least one diet **(Figure S4)**. The pattern of acylation changes on these proteins nearly follows the overall general trend of sugar-dependent hyper-acetylation and fat-dependent hypo-succinylation, except that ten of these acylation sites showed HFD-dependent hyperacetylation. Notably, six of the ten HFD-dependent hyper-acetylation sites were observed in proteins that regulate ammonia metabolism, including Cps1 (K287, K307, K458), GOT2 (K227, K309), and OTC (K231). This subset of amino acid metabolism proteins has been previously reported to be regulated by acylation (Nakagawa et al., 2009; Zhao et al., 2010).

### Discussion

In this system-wide, multi-omic study, we report widespread, global protein hyperacetylation resulting from chronic ingestion of dietary sugars, and widespread protein hypo-succinylation resulting from chronic HFD feeding. Accompanying protein-level changes support significant diet-induced remodeling of liver mitochondria metabolism, which was further reflected by altered metabolites. This quantitative analysis of diet-induced proteome, protein acylation, and metabolite remodeling together paints a comprehensive picture of the liver’s response to macronutrient excess.

Our metabolomic and proteomic quantification results reveal complementary aspects of the hepatic response to unique macronutrient stresses. Decreased pyruvate and lactate levels in mice fed HFD are likely explained by the increase in citrate, which is known to inhibit the key glycolytic enzyme PFK1 (Parmeggiani and Bowman, 1963; Randle et al., 1963). This increase in citrate levels was unlikely to be driven by differences in citrate synthase activity, which trended higher in all diets relative to controls **(Figure S5)**. Instead, the citrate accumulation observed in HFD-fed mice relative to glucose-fed mice appears to be linked to increased abundance of proteins involved in amino acid catabolism, an idea also supported by the clear decreases in abundance of nearly all amino acids measured. High levels of plasma citrate have been previously observed in patients with non-alcoholic fatty liver disease, where it was also found to promote the production of ROS (van de Wier et al., 2013), possibly due to chelation of ionic iron thereby allowing Fenton chemistry (Fenton, 1894). This is consistent with our data showing increased ROS in hepatocytes cultured with fatty acids (Softic et al., companion paper), and the decreases in ROS metabolizing/neutralizing enzymes such as SOD1 and glutathione metabolism proteins in mice fed HFD relative to mice fed excess glucose. The proteomic and metabolomic results therefore provide new mechanistic insight into the altered metabolic physiology of obesity.

The most salient finding from the acetylomic and succinylomic results is the dichotomy of sugar and fat dependent changes. Either sugar induced protein hyper-acetylation, whereas all of the HFD induced protein hypo-succinylation. Glucose-dependent hyperacetylation of MDH2 was previously reported in hepatocytes exposed to high concentrations of glucose (Zhao et al., 2010), but our study provides the first evidence of the global nature of this response *in vivo*. Whereas we found significant overlap in acetylation and succinylation sites, we observed very little crosstalk; acetylation sites that change in one diet rarely have alterations of succinylation in the same diet (see **Figure 4A, 4C, S1, and S3A).** This result is consistent with a non-enzymatic mechanism of mitochondrial protein acylation due to different routes of acetyl-CoA and succinyl-CoA production. These non-enzymatic modifiers may serve to block other forms of lysine modifications, such as ubiquitination and SUMOylation (Lumpkin et al., 2017).

Since the discovery of mitochondrial protein acylation, several studies have attempted to find site-specific functional effects of acylation using mutagenesis to change lysine into acyl-mimicking residues, such as lysine to glutamine, resulting in altered function. Experiments that introduce mutations to mimic acylation, using either recombinant protein *in vitro* or transfection of cells, reflect a state where all or most of the protein is acylated. However, several recent studies have shown that nearly all protein acylations are present at very low stoichiometry, with estimates of 1 in 1000 molecules acetylated (0.1%) (Baeza et al., 2014; Meyer et al., 2016; Weinert et al., 2017), raising concerns about the potential functional effects on protein activity. However, if protein acylation is activating, as is the case of tyrosine phosphorylation, then even a low % of modification could have a significant effect, for example by altering protein-protein interactions. Conversely, if protein acylation is inhibitory, the cumulative effects of these small enzyme inhibitions may result in major pathway-level activity changes. For example, if acetylation of ten proteins in a metabolic pathway are each reduced to 99% activity, then the overall flux through this pathway would be reduced to 90%. If instead those proteins were each reduced to 90% activity, then the total pathway activity could be as low as 35%. Thus, our data demonstrating modification of multiple proteins within specific metabolic pathways in response to changes in dietary composition could suggest that even low-level modifications might be contributing to regulation of metabolic fuel selection.

Previous studies reported that chronic HFD induces protein hyperacetylation (Alrob et al., 2014; Hirschey et al., 2011; Kendrick et al., 2011), which was not observed in the present study. In fact, recent research has concluded that acetyl-CoA from fat metabolism is the sole source of mitochondria protein hyperacetylation, and that fat drives histone acetylation (McDonnell et al., 2016; Pougovkina et al., 2014). However, previous experiments did not explicitly exclude sugar as a potential source of hyperacetylation and several experimental differences could account for the apparently disparate observations. First, compared to at least some experiments reported in previous studies, the present study used a different HFD with less sugar, i.e. ∼9% sucrose here, ∼29% sucrose in previous studies, as well as a different HFD composition. Second, previous studies administered HFD for at least 13 weeks, compared to this study where mice were fed HFD for only 10 weeks. Given that chronic HFD can induce elevated resting glucose (Pettersson et al., 2012) the hyper-acetylation observed from longer-term chronic HFD may actually result from diet-induced hyperglycemia, explaining the time dependence. Third, mice in previous studies were fasted before sacrifice whereas the mice used here were random-fed, and it is possible that acylation patterns can vary in fasted versus fed conditions. Consistent with this idea, lysine acetylation is known to increase in fasting in parallel with increases in circulating free fatty acids, and the fasting-induced increase in acetylation can be blocked by inhibition of lipolysis (Weinert et al., 2015). This shows that fatty acids can increase acetylation when they are a primary source of acetyl-CoA. Fourth, mice in some previous studies were sacrificed immediately at the start of the light cycle, whereas mice in this study were sacrificed slightly after the start of the light cycle. Sixth, in this study we used a different antibody mixture to enrich acetyl-lysine (Svinkina et al., 2015) than in some of the previous studies. Finally, differences in gut microbiota resulting from specific animal facilities are known to have strong impacts on metabolism (Ussar et al., 2015, 2016). We cannot explicitly rule out contributions from any of these variables. Nevertheless, under the conditions defined in the current study, we observed a clear dichotomy of effects of HF and sugar supplementation on protein acetylation and succinylation in animals sacrificed during ad-lib exposure to their respective diets.

Interpretation of the acyl-CoA and acyl-carnitine measurements in light of the acylation results provides further value to this resource. First, the HFD and HFD+F cohorts had the smallest changes in protein acetylation; however, these groups had the greatest accumulation of long-chain acyl-CoAs **(Fig. S2)**. Given the high concentrations of these numerous RACS, it is possible that mitochondrial protein lysine residues are acylated with longer-chain acyl groups that prevent acetylation and succinylation. Also, free CoA levels are elevated in every 10-week diet except those given excess dietary glucose, and the additional free-CoA may serve as a thiolate sink for acyl groups, again limiting lysine acylation. Finally, although acetyl-CoA and succinyl-CoA are needed for protein modification, their concentrations did not correlate with the observed patterns of protein acetylation or succinylation, indicating that acyl-CoA levels alone are not useful to predict protein acylation levels.

Liver plays a critical role in metabolic regulation by storing glucose as glycogen in the fed state and releasing glucose in the fasted state when lipids become the main fuel source. To achieve this, liver must regulate metabolic fuel selection across the spectrum of lipid, sugar and amino acid substrates, with metabolism of any of these fuels modifying the metabolism of the others (Randle et al., 1963; Hue and Taegtmeyer, 2009; Muoio, 2014). Our comprehensive dataset and analysis supports a dichotomous, nutrient-dependent model of protein acetylation and succinylation, defining a novel set of molecular responses to changes in diet composition that may contribute to modulation of mitochondrial protein function and metabolic fuel selection **(Figure 7A)**. Under our experimental conditions, acetyl-CoA from sugar metabolism uniquely drives mitochondrial protein acetylation. In contrast, in the setting of chronic HF feeding, acetyl-CoA derived from fatty-acid oxidation does not induce protein hyper-acetylation, even after 10 weeks of feeding. Instead HFD is associated with protein hypo-succinylation. HFD feeding also induces citrate accumulation, which is likely generated from fatty acid-derived acetyl CoA condensing with oxaloacetate derived from anaplerotic substrates, including amino acids, as suggested by their reduced levels in livers of HFD mice **(Figure 7B)**. HFD-dependent citrate accumulation is matched by pyruvate reduction, consistent with the classic model of fat-dependent inhibition of glycolysis. The divergent fates of acetyl-CoA and succinyl-CoA reported here may be an important regulatory mechanism contributing to loss of metabolic flexibility in conditions of nutrient overload and obesity.

**Figure 7:**
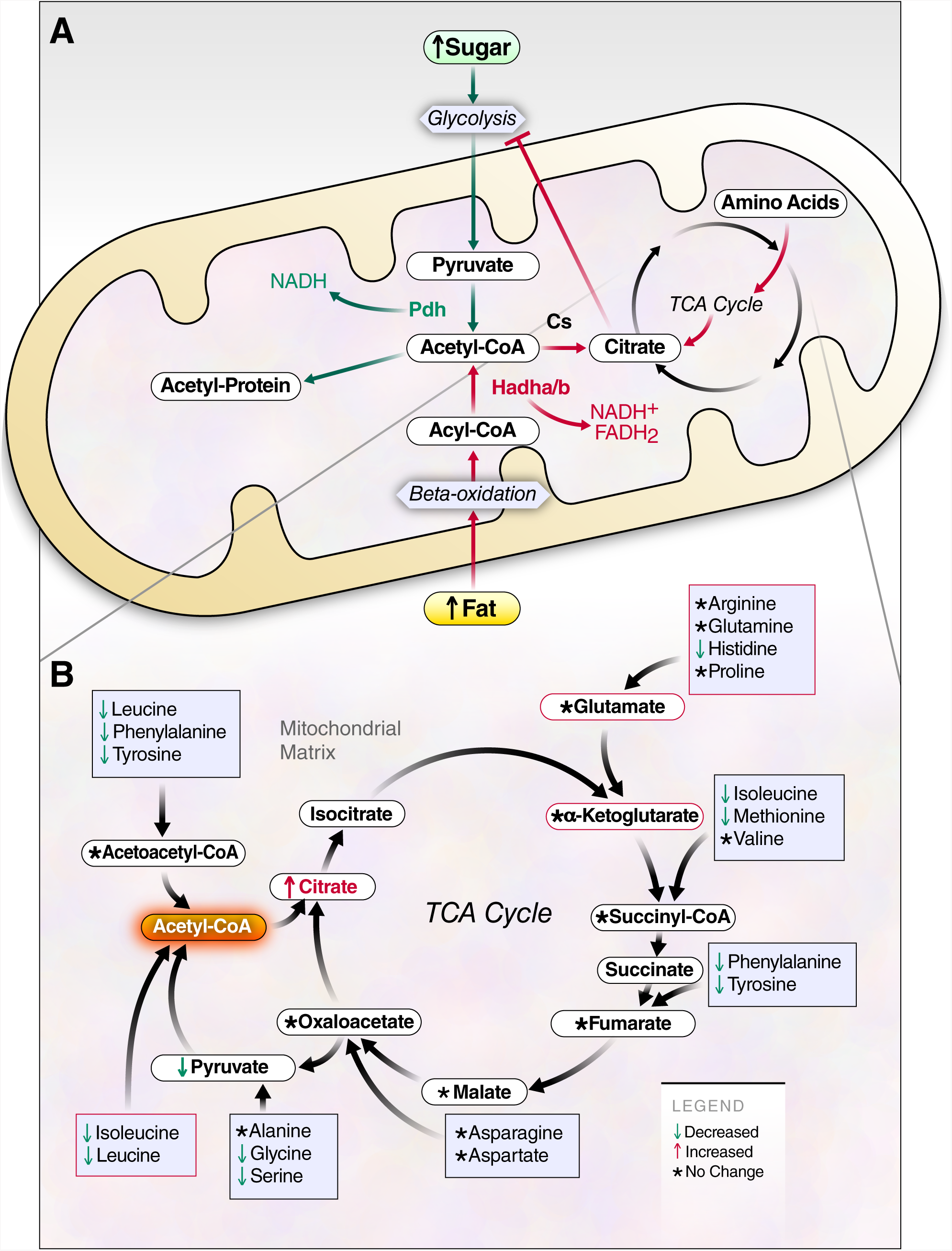
Models of dichotomous forces on mitochondrial functions and protein acylation resulting from excess dietary sugar or fat. **a,** Model describing the central dichotomy of acetyl-CoA production and fate resulting from either glucose or fat, drawn with green or red arrows, respectively. Although both glucose and fat are metabolized through the acetyl-CoA intermediate, acetyl-CoAs are not created equally; acetyl-CoA created from glycolysis results in protein hyperacetylation, but acetyl-CoA produced from fat metabolism does not cause hyperacetylation of protein. HFD is instead associated with citrate accumulation, which is known to inhibit glycolysis. **b,** Model describing the HFD-dependent production of citrate and anaplerosis through amino acid catabolism.

## Acknowledgments

We are grateful to John C. W. Carroll for help creating figures, and Dr. Matthew Hirschey for helpful discussions. This work was supported by funding from the NIH NIDDK (R24 DK085610 to E.V., C.R.K., B.S. and C.B.N.) and a NCRR shared instrumentation grant (1S10 OD016281 to B.W.G.). J.G.M. was supported by T32 AG000266, S.S. was supported by K12 HD000850, and N.B. was supported by a fellowship from the Glenn Foundation for Medical Research. This work was also supported by NIH grants R01 DK033201 (to C.R.K.) and the Joslin Diabetes Center DRC Animal Physiology and Bioinformatics Core facilities funded by P30 DK36836.

## Author Contributions

Conceptualization, J.G.M., S.S., N.B., M.J.R., E.V., B.W.G., C.B.N., C.R.K., and B.S.; Methodology, J.G.M., S.S., N.B., B.W.G., C.B.N., and C.R.K., Software, J.G.M., Formal Analysis, J.G.M., Investigation, J.G.M. and S.S., Resources, B.S., B.W.G., C.B.N., C.R.K., and S.S.; Data Curation, J.G.M.; Writing – Original Draft, J.G.M., Writing – Reviewing and Editing, J.G.M., S.S., N.B., B.W.B., C.B.N., C.R.K., and B.S.; Visualization, J.G.M.; Supervision, E.V., C.B.N., C.R.K., B.W.G., B.S.; Project Administration, E.V., B.W.G., C.B.N., C.R.K., B.S.; Funding Acquisition, E.V., C.B.N., C.R.K., and B.W.G.

## Declaration of Interests

The authors declare no competing financial interests.

## STAR Methods

### CONTACT FOR REAGENT AND RESOURCE SHARING

Birgit Schilling

### EXPERIMENTAL MODEL AND SUBJECT DETAILS

#### Animals and diets

All protocols were approved by the IACUC of the Joslin Diabetes Center and were in accordance with NIH guidelines. Mice were housed at 20-22°C on a 12 h light/dark cycle with ad libitum access to food and water. C57Bl6, male mice at 6 weeks of age were purchased from Jackson Laboratory and fed either chow diet (Mouse Diet 9F, PharmaServ) or HFD (Research diets, D12492) for 10 weeks. Caloric composition of chow diet consisted of 23% protein, 21.6% fat and 55.4% carbohydrates, while HFD had 20% protein, 60% fat and 20% carbohydrates. Mice were watered with either tap water or 30% (w/v) fructose or 30% glucose solution in water.

### METHOD DETAILS

Mice were weighed and their food intake was recorded once per week. Mice were sacrificed from 8 to 11am, and one mouse from each cage, i.e. dietary group, was utilized before sacrificing the next mouse in the same cage. This was repeated until all four mice per cage were sacrificed. Serum parameters were determined by the Joslin Diabetes and Endocrinology Research Center assay core using commercial colorimetric assays for beta-hydroxybutyrate and free fatty acids as previously reported (Softic et al., 2017).

#### Mitochondria protein digestion and acetylated peptide enrichment

Mitochondria protein was quantified using the BCA assay, and mitochondria containing 1 mg of protein were lysed with addition of 100 μL of 8M Urea, 10 μL of 10% maltoside, 10 μL of 1M TEAB buffer (pH 8.5), and water to a final volume of 200 μL Proteins were reduced for 30 minutes at 37 ° Celsius by addition of DTT to a final concentration of 4.5 mM. Reduced protein solutions were then cooled to room temperature and alkylated by the addition of iodoacetamide to a final concentration of 10 mM for 30 min at room temperature. The alkylated protein solution was then diluted to 1 mL final volume using 50 mM TEAB, pH 8.5, and trypsin was added at a ratio of 1 part trypsin per 50 parts mitochondrial protein (wt:wt, 20 μg per sample). Trypsin digestion proceeded overnight for 18 hours at 37° C. Digestion was quenched by addition of formic acid to a final concentration of 1%, and insoluble material was precipitated by centrifugation at 1,800 relative centrifugal force for 15 min at room temperature. Soluble peptides were then desalted using Waters’ Oasis HLB Vacuum cartridges (30 mg sorbent, 1 mL volume). Peptides were eluted with 1.2 mL of 80% acetonitrile (ACN) / 0.2% formic acid (FA) /19.8% water, and dried to completion using a vacuum centrifugal concentrator. Desalted peptides were then resuspended in 1.4 mL of IAP buffer (50mM MOPS, 10 mM Na2PO4, 50 mM NaCl) for immunoprecipitation. Acetylated peptides were enriched using anti-acetyl lysine antibody-bead conjugate (PTMScan #13416, Cell Signaling Technologies) using 10 μL of beads per sample (1/4 of the manufacturer-supplied tube per sample, approximately 62.5 μg of conjugated antibody). Immunoprecipitation proceeded overnight at 4° C with rocking. The following day, supernatant containing unbound peptides was removed and saved for subsequent anti-succinyl-lysine IP. The beads were washed twice with 1 mL of ice-cold IAP buffer, three times with ice-cold water (Burdick and Jackson HPLC-grade), and then eluted using 100 μL of 0.15% trifluoracetic acid (55 μL followed by a second elution using 45 μL Unbound peptides from the first IP were then used as input for a second IP using the same method but anti-succinyl lysine antibody-bead conjugate (PTMScan #13764, Cell Signaling Technologies). Peptides eluted from the enrichment were desalted directly using C18 reversed phase StageTips (two eighteen-gauge disks per StageTip), and resuspended in 7 μL of 0.2% formic acid for mass spectrometry analysis.

#### Nanoflow liquid chromatography – tandem mass spectrometry

Peptide separations were carried out using mobile phase A consisting of 97.95% water/0.05% FA/2% acetonitrile, and mobile phase B consisting of 98% acetonitrile/1.95% water/0.05% FA. Samples were loaded onto the first of two sequential C18 column chips (75 μm x 15 cm ChromXP C18-CL chip, 3 μm particles, 300 Å) using an Eksigent cHiPLC system for 30 min at a flow of 0.6 μL per min of mobile phase A. Separation was performed over two 75 μm x 15 cm ChromXP C18-CL analytical chip (30 cm total length) using a gradient from 5% to 40% mobile phase B over 80 minutes. The column chip was washed by increasing to 80% B over 5 minutes that was maintained for 8 minutes, followed by a return to 5% mobile phase B over two minutes that was maintained for 25 minutes to re-equilibrate the column. Eluting peptides were directly electrosprayed into a TripleTOF 5600 mass spectrometer (SCIEX) and analyzed by SWATH data-independent acquisition (DIA). Every SWATH cycle consisted of a 250 ms precursor ion scan from 400-1,250 m/z followed by fragmentation of all ions between 400-1,200 m/z using 64 variable width precursor isolation windows. The SWATH window definitions used for enrichments of acetylated peptides were determined based on the frequency of acetylated peptide identifications in each m/z region using previous data-dependent acquisitions for 42 ms each, resulting in a total cycle time of approximately 3 sec. Non-enriched peptides were also analyzed by SWATH to determine protein-level changes. The SWATH window definitions used to collect data for protein-level changes were based on the cross-lab SWATH study (Collins et al., 2017). Fragment ion spectra were collected from 100-2,000 m/z.

#### Data analysis

Data for protein-level analysis was analyzed using Spectronaut (Bruderer et al., 2015) with our in-house spectral library from DDA runs of pooled protein samples. DIA data from acetyl-peptide enrichment was processed using PIQED (Meyer et al., 2017), which can automatically identify and quantify site-level PTMs using only DIA-MS data, using settings described in the original manuscript. PIQED strings together several tools to accomplish this, including mapDIA (Teo et al., 2015), TPP tools (Deutsch et al., 2015), and DIA-Umpire (Tsou et al., 2015). PIQED analysis used protein-level quantities determined using Spectronaut for correction of site-level changes by protein-level changes. Additional analysis was carried out using custom scripts written in R (R Development Core Team, 2008). Functional analysis of protein changes was performed using ClueGO within Cytoscape (Bindea et al., 2009; Shannon et al., 2003).

##### Metabolite quantification

50 mg of previously frozen liver tissue was homogenized in 1 ml of 50% aqueous ACN, containing 0.3% FA. Liver amino acids, acyl CoAs and organic acids were analyzed using stable isotope dilution techniques. Acyl carnitines were measured by LC-MS/MS as described previously (White et al., 2016).

## QUANTIFICATION AND STATISTICAL ANALYSIS

### Statistical analyses

Significant differences for protein levels were computed using student’s t-tests within spectronaut, and significant differences between acetylation sites and succinylation sites were computed using a Bayesian model within mapDIA (Teo et al., 2015) as part of the PIQED software tool and workflow. Significant protein level changes were defined as log2(FC) >0.58 and q-value < 0.01, and significant site-level changes were defined as log2(FC)>1 and q-value < 0.01. On heatmaps, sites not meeting significance are plotted in grey or white. Fischer exact testing was used to determine functional term enrichment analysis within the ClueGO plugin of cytoscape. From each cohort, we collected data for 5 biological replicates, but due to mass spectrometer software errors, we lost data the fifth replicate of the acetylation enrichment from 10-week HFD+G. Among the succinylation replicates, we lost data for the fifth replicates of the succinylation enrichments from: 2-week F, 2-week G, 10-week F, 10-week HFD, and 10-week HFD+G. From the 2-week control succinylation set, we lost data from two samples, leaving 3 replicates for that group. Therefore, all cohorts had at least 3 biological replicates, but most cohorts had 5 replicates.

## DATA AND SOFTWARE AVAILABILITY

The Massive repository (ftp://massive.ucsd.edu or ftp://MSV000081902@massive.ucsd.edu username: MSV000081902, password: dietstudy) contains all the raw mass spectrometry data files used for quantification of proteins, protein acetylation, and protein succinylation, as well as the raw files used to generate a sample-specific protein-level spectral library for quantification using Spectronaut. The massive repository also contains several supplemental files, including: “ptmProphet-acetyl.ptm.pep.xml” and “ptmProphet-succinyl.ptm.pep.xml” containing the Peptideprophet-, iProphet-, and PTMprophet-filtered identifications produced using PIQED.

“RAW_MS_descriptions.txt” containing a legend describing MS file naming, “aK_SkylineReport.csv” and “sK_SkylineReport.csv” containing fragment-level peak areas for acetylated and succinylated peptides, respectively, “mapDIA_Input_aK.txt” and “mapDIA_Input_sK.txt” containing the inputs for mapDIA statistical testing for acetyl and succinyl data, respectively, “mapDIA_analysis_output_aK.txt” and “mapDIA_analysis_output_sK.txt” containing the site-level fold changes and FDRs computed by mapDIA for acetyl and succinyl data, respectively, “dia_umpire_se_params.txt” containing the settings used by PIQED for running DIA-Umpire “20150810.mouse.cc.iRT.fasta” protein database used by PIQED for MS-GF+ search and “20150810.mouse.cc.iRT_DECOY.fasta” used by PIQED for Comet and X!Tandem searches.

All the processed acetylation and succinylation data is available on the Panorama: https://panoramaweb.org/labkey/project/Schilling/Meyer_acylomes_diets/begin.view?

## Supplementary Files

Five supplementary figures are contained in the file “supp_figs_final.pdf”. Details of protein identifications used for quantification are given in **Table S1**. Details of acetylated peptide identifications are given in **Table S2**, and succinylated peptide identifications are given in **Table S3**. **Table S4** gives the quantitative results for proteins. **Table S5** gives quantitative results for acetylation sites, and **Table S6** gives quantitative results for succinylation Sites.

## References

Alrob, O.A., Sankaralingam,S., Ma, C., Wagg, C.S., Fillmore, N., Jaswal, J.S., Sack, M.N., Lehner, R., Gupta, M.P., Michelakis, E.D., et al. (2014). Obesity-induced lysine acetylation increases cardiac fatty acid oxidation and impairs insulin signalling. Cardiovasc. Res. 103, 485–497.

Baeza, J., Dowell, J.A., Smallegan, M.J., Fan, J., Amador-Noguez, D., Khan, Z., and Denu, J.M. (2014). Stoichiometry of site-specific lysine acetylation in an entire proteome. J. Biol. Chem.

Baeza, J., Smallegan, M.J., and Denu, J.M. (2015). Site-Specific Reactivity of Nonenzymatic Lysine Acetylation. ACS Chem. Biol. 10, 122–128.

Bharathi, S.S., Zhang, Y., Mohsen, A.-W., Uppala, R., Balasubramani, M., Schreiber, E., Uechi, G., Beck, M.E., Rardin, M.J., Vockley, J., et al. (2013). Sirtuin 3 (SIRT3) Protein Regulates Long-chain Acyl-CoA Dehydrogenase by Deacetylating Conserved Lysines Near the Active Site. J. Biol. Chem. 288, 33837–33847.

Bheda, P., Jing, H., Wolberger, C., and Lin, H. (2016). The Substrate Specificity of Sirtuins. Annu. Rev. Biochem. 85, 405–429.

Bindea, G., Mlecnik, B., Hackl, H., Charoentong, P., Tosolini, M., Kirilovsky, A., Fridman, W.-H., Pagès, F., Trajanoski, Z., and Galon, J. (2009). ClueGO: a Cytoscape plug-in to decipher functionally grouped gene ontology and pathway annotation networks. Bioinformatics 25, 1091–1093.

Bricambert, J., Miranda, J., Benhamed, F., Girard, J., Postic, C., and Dentin, R. (2010). Salt-inducible kinase 2 links transcriptional coactivator p300 phosphorylation to the prevention of ChREBP-dependent hepatic steatosis in mice. J. Clin. Invest. 120, 4316–4331.

Bruderer, R., Bernhardt, O.M., Gandhi, T., Miladinović, S.M., Cheng, L.-Y., Messner, S., Ehrenberger, T., Zanotelli, V., Butscheid, Y., Escher, C., et al. (2015). Extending the Limits of Quantitative Proteome Profiling with Data-Independent Acquisition and Application to Acetaminophen-Treated Three-Dimensional Liver Microtissues. Mol. Cell. Proteomics MCP 14, 1400–1410.

Collins, B.C., Hunter, C.L., Liu, Y., Schilling, B., Rosenberger, G., Bader, S.L., Chan, D.W., Gibson, B.W., Gingras, A.-C., Held, J.M., et al. (2017). Multi-laboratory assessment of reproducibility, qualitative and quantitative performance of SWATH-mass spectrometry. Nat. Commun. 8, 291.

Deutsch, E.W., Mendoza, L., Shteynberg, D., Slagel, J., Sun, Z., and Moritz, R.L. (2015). Trans-Proteomic Pipeline, a standardized data processing pipeline for large-scale reproducible proteomics informatics. PROTEOMICS – Clin. Appl. 9, 745–754.

Fenton, H.J.H. (1894). LXXIII.-Oxidation of tartaric acid in presence of iron. J. Chem. Soc. Trans. 65, 899–910.

Frayn, K.N., and Kingman, S.M. (1995). Dietary sugars and lipid metabolism in humans. Am. J. Clin. Nutr. 62, 250S–261S.

Galgani, J.E., Moro, C., and Ravussin, E. (2008). Metabolic flexibility and insulin resistance. Am. J. Physiol.-Endocrinol. Metab. 295, E1009–E1017.

Ghanta, S., Grossmann, R.E., and Brenner, C. (2013). Mitochondrial protein acetylation as a cell-intrinsic, evolutionary driver of fat storage: Chemical and metabolic logic of acetyl-lysine modifications. Crit. Rev. Biochem. Mol. Biol. 48, 561–574.

Gillet, L.C., Navarro, P., Tate, S., Röst, H., Selevsek, N., Reiter, L., Bonner, R., and Aebersold, R. (2012). Targeted Data Extraction of the MS/MS Spectra Generated by Data-independent Acquisition: A New Concept for Consistent and Accurate Proteome Analysis. Mol. Cell. Proteomics 11.

Hebert, A.S., Dittenhafer-Reed, K.E., Yu, W., Bailey, D.J., Selen, E.S., Boersma, M.D., Carson, J.J., Tonelli, M., Balloon, A.J., Higbee, A.J., et al. (2013). Calorie Restriction and SIRT3 Trigger Global Reprogramming of the Mitochondrial Protein Acetylome. Mol. Cell 49, 186–199.

Hirschey, M., Shimazu, T., Jing, E., Grueter, C., Collins, A., Aouizerat, B., Stan??kov?, A., Goetzman, E., Lam, M., Schwer, B., et al. (2011). SIRT3 Deficiency and Mitochondrial Protein Hyperacetylation Accelerate the Development of the Metabolic Syndrome. Mol. Cell 44, 177–190.

Hirschey, M.D., Shimazu, T., Goetzman, E., Jing, E., Schwer, B., Lombard, D.B., Grueter, C.A., Harris, C., Biddinger, S., Ilkayeva, O.R., et al. (2010). SIRT3 regulates mitochondrial fatty-acid oxidation by reversible enzyme deacetylation. Nature 464, 121–125.

Hue, L., and Taegtmeyer, H. (2009). The Randle cycle revisited: a new head for an old hat. Am. J. Physiol. - Endocrinol. Metab. 297, E578.

James, A.M., Hoogewijs, K., Logan, A., Hall, A.R., Ding, S., Fearnley, I.M., and Murphy, M.P. (2017). Non-enzymatic N-acetylation of Lysine Residues by AcetylCoA Often Occurs via a Proximal S-acetylated Thiol Intermediate Sensitive to Glyoxalase II. Cell Rep. 18, 2105–2112.

Kendrick, A.A., Choudhury, M., Rahman, S.M., McCurdy, C.E., Friederich, M., Van Hove, J.L.K., Watson, P.A., Birdsey, N., Bao, J., Gius, D., et al. (2011). Fatty liver is associated with reduced SIRT3 activity and mitochondrial protein hyperacetylation. Biochem. J. 433, 505–514.

Levin, E., Lopez-Martinez, G., Fane, B., and Davidowitz, G. (2017). Hawkmoths use nectar sugar to reduce oxidative damage from flight. Science 355, 733.

Li, F., He, X., Ye, D., Lin, Y., Yu, H., Yao, C., Huang, L., Zhang, J., Wang, F., Xu, S., et al. (2015). NADP+-IDH Mutations Promote Hypersuccinylation that Impairs Mitochondria Respiration and Induces Apoptosis Resistance. Mol. Cell 60, 661–675.

Lumpkin, R.J., Gu, H., Zhu, Y., Leonard, M., Ahmad, A.S., Clauser, K.R., Meyer, J.G., Bennett, E.J., and Komives, E.A. (2017). Site-specific identification and quantitation of endogenous SUMO modifications under native conditions. Nat. Commun. 8, 1171.

McDonnell, E., Crown, S.B., Fox, D.B., Kitir, B., Ilkayeva, O.R., Olsen, C.A., Grimsrud, P.A., and Hirschey, M.D. (2016). Lipids Reprogram Metabolism to Become a Major Carbon Source for Histone Acetylation. Cell Rep. 17, 1463–1472.

Merrell, M.D., and Cherrington, N.J. (2011). Drug metabolism alterations in nonalcoholic fatty liver disease. Drug Metab. Rev. 43, 317–334.

Meyer, J.G., D’Souza, A.K., Sorensen, D.J., Rardin, M.J., Wolfe, A.J., Gibson, B.W., and Schilling, B. (2016). Quantification of Lysine Acetylation and Succinylation Stoichiometry in Proteins Using Mass Spectrometric Data-Independent Acquisitions (SWATH). J. Am. Soc. Mass Spectrom. 27, 1758–1771.

Meyer, J.G., Mukkamalla, S., Steen, H., Nesvizhskii, A.I., Gibson, B.W., and Schilling, B. (2017). PIQED: automated identification and quantification of protein modifications from DIA-MS data. Nat Meth 14, 646–647.

Muoio, D.M. (2014). Metabolic Inflexibility: When Mitochondrial Indecision Leads to Metabolic Gridlock. Cell 159, 1253–1262.

Nakagawa, T., Lomb, D.J., Haigis, M.C., and Guarente, L. (2009). SIRT5 Deacetylates Carbamoyl Phosphate Synthetase 1 and Regulates the Urea Cycle. Cell 137, 560–570.

Newman, J.C., Covarrubias, A.J., Zhao, M., Yu, X., Gut, P., Ng, C.-P., Huang, Y., Haldar, S., and Verdin, E. (2017). Ketogenic Diet Reduces Midlife Mortality and Improves Memory in Aging Mice. Cell Metab. 26, 547–557.e8.

Newsholme, E.A., Sugden, P.H., and Williams, T. (1977). Effect of citrate on the activities of 6-phosphofructokinase from nervous and muscle tissues from different animals and its relationships to the regulation of glycolysis. Biochem. J. 166, 123.

Nishida, Y., Rardin, M.J., Carrico, C., He, W., Sahu, A.K., Gut, P., Najjar, R., Fitch, M., Hellerstein, M., Gibson, B.W., et al. (2015). SIRT5 Regulates both Cytosolic and Mitochondrial Protein Malonylation with Glycolysis as a Major Target. Mol. Cell 59, 321–332.

Park, E.J., Lee, J.H., Yu, G.-Y., He, G., Ali, S.R., Holzer, R.G., Österreicher, C.H., Takahashi, H., and Karin, M. (2010). Dietary and genetic obesity promote liver inflammation and tumorigenesis by enhancing IL-6 and TNF expression. Cell 140, 197–208.

Park, J., Chen, Y., Tishkoff, D.X., Peng, C., Tan, M., Dai, L., Xie, Z., Zhang, Y., Zwaans, B.M.M., Skinner, M.E., et al. (2013). SIRT5-Mediated Lysine Desuccinylation Impacts Diverse Metabolic Pathways. Mol. Cell 50, 919–930.

Parmeggiani, A., and Bowman, R.H. (1963). Regulation of phosphofructokinase activity by citrate in normal and diabetic muscle. Biochem. Biophys. Res. Commun. 12, 268–273.

Pettersson, U.S., Waldén, T.B., Carlsson, P.-O., Jansson, L., and Phillipson, M. (2012). Female Mice are Protected against High-Fat Diet Induced Metabolic Syndrome and Increase the Regulatory T Cell Population in Adipose Tissue. PLOS ONE 7, e46057.

Picklo, M.J. (2008). Ethanol intoxication increases hepatic N-lysyl protein acetylation. Biochem. Biophys. Res. Commun. 376, 615–619.

Pougovkina, O., te Brinke, H., Ofman, R., van Cruchten, A.G., Kulik, W., Wanders, R.J.A., Houten, S.M., and de Boer, V.C.J. (2014). Mitochondrial protein acetylation is driven by acetyl-CoA from fatty acid oxidation. Hum. Mol. Genet. 23, 3513–3522.

R Development Core Team (2008). R: A language and environment for statistical computing (Vienna, Austria: R Foundation for Statistical Computing).

Randle, P.J., Garland, P.B., Hales, C.N., and Newsholme, E.A. (1963). THE GLUCOSE FATTY-ACID CYCLE ITS ROLE IN INSULIN SENSITIVITY AND THE METABOLIC DISTURBANCES OF DIABETES MELLITUS. The Lancet 281, 785–789.

Rardin, M.J., He, W., Nishida, Y., Newman, J.C., Carrico, C., Danielson, S.R., Guo, A., Gut, P., Sahu, A.K., Li, B., et al. (2013a). SIRT5 regulates the mitochondrial lysine succinylome and metabolic networks. Cell Metab. 18, 920–933.

Rardin, M.J., Newman, J.C., Held, J.M., Cusack, M.P., Sorensen, D.J., Li, B., Schilling, B., Mooney, S.D., Kahn, C.R., Verdin, E., et al. (2013b). Label-free quantitative proteomics of the lysine acetylome in mitochondria identifies substrates of SIRT3 in metabolic pathways. Proc. Natl. Acad. Sci. 110, 6601–6606.

Robles, M.S., Humphrey, S.J., and Mann, M. (2017). Phosphorylation Is a Central Mechanism for Circadian Control of Metabolism and Physiology. Cell Metab. 25, 118–127.

Sadhukhan, S., Liu, X., Ryu, D., Nelson, O.D., Stupinski, J.A., Li, Z., Chen, W., Zhang, S., Weiss, R.S., Locasale, J.W., et al. (2016). Metabolomics-assisted proteomics identifies succinylation and SIRT5 as important regulators of cardiac function. Proc. Natl. Acad. Sci. 113, 4320–4325.

Saxholm, H.J.K., Pestana, A., O’Connor, L., Sattler, C.A., and Pitot, H.C. (1982). Protein acetylation. Mol. Cell. Biochem. 46, 129–153.

Schwer, B., Eckersdorff, M., Li, Y., Silva, J.C., Fermin, D., Kurtev, M.V., Giallourakis, C., Comb, M.J., Alt, F.W., and Lombard, D.B. (2009). Calorie restriction alters mitochondrial protein acetylation. Aging Cell 8, 604–606.

Shannon, P., Markiel, A., Ozier, O., Baliga, N.S., Wang, J.T., Ramage, D., Amin, N., Schwikowski, B., and Ideker, T. (2003). Cytoscape: a software environment for integrated models of biomolecular interaction networks. Genome Res. 13, 2498–2504.

Softic, S., Gupta, M.K., Wang, G.-X., Fujisaka, S., O’Neill, B.T., Rao, T.N., Willoughby, J., Harbison, C., Fitzgerald, K., Ilkayeva, O., et al. (2017). Divergent effects of glucose and fructose on hepatic lipogenesis and insulin signaling. J. Clin. Invest. 127, 4059–4074.

Sol, E.M., Wagner, S.A., Weinert, B.T., Kumar, A., Kim, H.-S., Deng, C.-X., and Choudhary, C. (2012). Proteomic Investigations of Lysine Acetylation Identify Diverse Substrates of Mitochondrial Deacetylase Sirt3. PLOS ONE 7, e50545.

Svinkina, T., Gu, H., Silva, J.C., Mertins, P., Qiao, J., Fereshetian, S., Jaffe, J.D., Kuhn, E., Udeshi, N.D., and Carr, S.A. (2015). Deep, Quantitative Coverage of the Lysine Acetylome Using Novel Anti-acetyl-lysine Antibodies and an Optimized Proteomic Workflow. Mol. Cell. Proteomics 14, 2429–2440.

Teo, G., Kim, S., Tsou, C.-C., Collins, B., Gingras, A.-C., Nesvizhskii, A.I., and Choi,H. (2015). mapDIA: Preprocessing and statistical analysis of quantitative proteomics data from data independent acquisition mass spectrometry. J. Proteomics 129, 108–120.

Thomson, S.J., Askari, A., and Bishop-Bailey, D. (2012). Anti-Inflammatory Effects of Epoxyeicosatrienoic Acids. Int. J. Vasc. Med. 2012, 605101.

Tsou, C.-C., Avtonomov, D., Larsen, B., Tucholska, M., Choi, H., Gingras, A.-C., and Nesvizhskii, A.I. (2015). DIA-Umpire: comprehensive computational framework for data-independent acquisition proteomics. Nat Meth 12, 258–264.

Ussar, S., Griffin, N.W., Bezy, O., Fujisaka, S., Vienberg, S., Softic, S., Deng, L., Bry, L., Gordon, J.I., and Kahn, C.R. (2015). Interactions between Gut Microbiota, Host Genetics and Diet Modulate the Predisposition to Obesity and Metabolic Syndrome. Cell Metab. 22, 516–530.

Ussar, S., Fujisaka, S., and Kahn, C.R. (2016). Interactions between host genetics and gut microbiome in diabetes and metabolic syndrome. Mol. Metab. 5, 795–803.

Wagner, G.R., and Hirschey, M.D. (2014). Nonenzymatic Protein Acylation as a Carbon Stress Regulated by Sirtuin Deacylases. Mol. Cell 54, 5–16.

Wagner, G.R., Bhatt, D.P., O’Connell, T.M., Thompson, J.W., Dubois, L.G., Backos, D.S., Yang, H., Mitchell, G.A., Ilkayeva, O.R., Stevens, R.D., et al. (2017). A Class of Reactive Acyl-CoA Species Reveals the Non-enzymatic Origins of Protein Acylation. Cell Metab. 25, 823–837.e8.

Weinert, B.T., Schölz, C., Wagner, S.A., Iesmantavicius, V., Su, D., Daniel, J.A., and Choudhary, C. (2013). Lysine Succinylation Is a Frequently Occurring Modification in Prokaryotes and Eukaryotes and Extensively Overlaps with Acetylation. Cell Rep. 4, 842–851.

Weinert, B.T., Moustafa, T., Iesmantavicius, V., Zechner, R., and Choudhary, C. (2015). Analysis of acetylation stoichiometry suggests that SIRT3 repairs nonenzymatic acetylation lesions. EMBO J. 34, 2620–2632.

Weinert, B.T., Satpathy, S., Hansen, B.K., Lyon, D., Jensen, L.J., and Choudhary, C. (2017). Accurate Quantification of Site-specific Acetylation Stoichiometry Reveals the Impact of Sirtuin Deacetylase CobB on the E. coli Acetylome. Mol. Cell. Proteomics 16, 759–769.

White, P.J., Lapworth, A.L., An, J., Wang, L., McGarrah, R.W., Stevens, R.D., Ilkayeva, O., George, T., Muehlbauer, M.J., Bain, J.R., et al. (2016). Branched-chain amino acid restriction in Zucker-fatty rats improves muscle insulin sensitivity by enhancing efficiency of fatty acid oxidation and acyl-glycine export. Mol. Metab. 5, 538–551.

van de Wier, B., Balk, J.M., Haenen, G.R.M.M., Giamouridis, D., Bakker, J.A., Bast, B.C., den Hartog, G.J.M., Koek, G.H., and Bast, A. (2013). Elevated citrate levels in non-alcoholic fatty liver disease: The potential of citrate to promote radical production. FEBS Lett. 587, 2461–2466.

Zhang, Z., Tan, M., Xie, Z., Dai, L., Chen, Y., and Zhao, Y. (2011). Identification of lysine succinylation as a new post-translational modification. Nat Chem Biol 7, 58–63.

Zhao, S., Xu, W., Jiang, W., Yu, W., Lin, Y., Zhang, T., Yao, J., Zhou, L., Zeng, Y., Li, H., et al. (2010). Regulation of Cellular Metabolism by Protein Lysine Acetylation. Science 327, 1000–1004.

Zhu, D., Hou, L., Hu, B., Zhao, H., Sun, J., Wang, J., and Meng, X. (2016). Crosstalk among proteome, acetylome and succinylome in colon cancer HCT116 cell treated with sodium dichloroacetate. Sci. Rep. 6.

